# Simple but Robust Improvement in Multivoxel Pattern Classification

**DOI:** 10.1101/326389

**Authors:** Sangil Lee, Joseph W. Kable

## Abstract

Multivoxel pattern analysis (MVPA) typically begins with the estimation of single trial activation levels, and several studies have examined how different procedures for estimating single trial activity affect the ultimate classification accuracy of MVPA. Here we show that the currently preferred estimation procedures impart spurious shifts in run-level means that cause the estimated activities to be misaligned across runs. These shifts are caused by positive correlations between the means of different category activity estimates within the same scanner run. In other words, if the mean of the estimates for one type of trials is high (low) in a given scanner run, then the mean of the other type of trials is also high (low) for that same scanner run, and the mean across all trials therefore shifts from run to run. Simulations show that these correlations are unavoidable whenever there is a need to deconvolve overlapping trial activities in the presence of noise. We show that subtracting the mean across all trials of a run from all the estimates within that run (i.e., run-level mean centering of estimates), by cancelling out these mean shifts, leads to robust and significant improvements in MVPA classification accuracy. These improvements are seen in both simulated and real data across a wide variety of situations and can provide significant direct benefits with no computational cost. However, we also point out that there could be cases when mean activations are expected to shift across runs and that run-level mean centering could be detrimental in some of these cases (e.g., different proportion of trial types between different runs).

---

Multivoxel pattern analysis (MVPA) has become an indispensable tool in fMRI research, enabling the classification of different types of tasks, stimuli, and/or mental states based on fMRI data (Haynes & Rees, 2006; Norman et al., 2006; Pereira, Mitchell, & Botvinick, 2009). Numerous studies have examined different ways to improve the classification power of MVPA through feature selection/data reduction (e.g., Chang et al., 2015; De Martino et al., 2008) or the use of different classifiers (e.g., Misaki et al., 2010). Several of these studies have focused on how to obtain single trial activation estimates *(i.e.,* the level of BOLD activity on each trial).

Many different methods have been proposed for estimating single trial activity, and two methods, in particular, have been studied in detail: 1) beta-series regression or LS-A (Least Squares-All; Rissman et al., 2004), and 2) LS-S (Least Squares - Separate; Turner, 2010). LS-A includes the simulated BOLD response of each trial as a separate regressor in one general linear model (GLM) and uses the coefficients for these regressors as the single trial activity estimates. LS-S, on the other hand, involves running a separate regression to estimate the activity on each trial, with each regression including one regressor modeling the trial of interest and one nuisance regressor modeling the response on all other trials. Mumford et al. (2012) have shown that LS-A is prone to multicollinearity that may weaken classification performance, and that LS-S provides better classification accuracies than several other methods, including ridge regression, partial least squares, and support vector regression. Turner et al. (2012) have shown that including a separate nuisance regressor for each different category of trial type further improves the performance of LS-S. On the other hand, Abdulrahman & Henson (2016) demonstrated that when there is relatively little noise, the regularization effect of the nuisance regressor(s) can become harmful and that LS-A may perform better under these conditions. Both methods, however, were shown to result in correlations between the estimated activities of neighboring trials (Mumford et al., 2014). With LS-A, there was a negative correlation between neighboring trials, while with LS-S there was a positive correlation between neighbors.

Here we explore a related but distinct issue. In simulations where the true mean activities for different trial types are known to be independent from each other and stable from run to run, we find that both LS-A and LS-S return estimated mean activities that are correlated across different trial types within a run. In other words, if the mean of the estimated activity levels for one category of trials in a run is high (low), then the mean of the estimated activity levels for the other category of trials in that same run is also high (low). Accordingly, LS-A and LS-S introduce spurious shifts in the estimated mean of all trial types from scanner run to scanner run. We observe that LS-A and LS-S estimates have these features whenever there is a need to estimating overlapping trial activities in the presence of any source of noise. Spurious shifts in the estimated mean of all trials of a run across different scanner runs could reduce classification accuracy, as many classification techniques seek to estimate a stable boundary between different trial types. However, a simple way to counteract this would be to mean center estimates within a run, subtracting out the mean of all the estimates in a run from each of the trial-wise estimates within that run. We show that such run-wise mean centering significantly improves classification accuracies in both simulated and real data. Importantly, however, we also explore circumstances under which there are expected to be true signal-related shifts in the mean activity of all trials of a run between different runs, such as when there have been shifts in a subjects’ baseline attentional or physiological state or when runs have different proportions of different trial types. We show the effect of run-wise mean centering in such scenarios and offer boundary conditions for its usefulness.

## Methods

### Data and Analysis

#### Simulated Data

Our simulation procedure largely followed that of Mumford et al. (2012). Each simulated scanner run contained two different trial types: type A and type B. The true trial activation levels were simulated as the following:

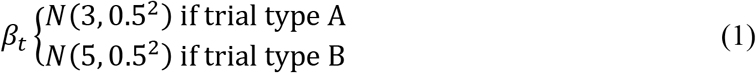

The ISI was drawn from a uniform distribution of *U(ISImin, ISImax)*. This is because the uniform distribution gives us control over the mean while holding the variance constant. Though not reported in detail here, we have also used exponentially distributed ISIs with means that match the uniform distributions and in all cases we observe similar results. Finally, a voxel’s signal was generated by 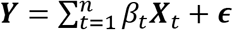 where ***X***_*t*_ is the expected BOLD activity for trial *t* (i.e., same as an LS-A regressor, created by convolving a simulated boxcar of neural activity with canonical HRF), and ***∊*** ~ *N*(0, *σ*^2^***V***) where ***V*** is an AR(1) correlation matrix. The value of auto-correlation was matched to Mumford et al. (2012) as 0.12 between TRs for all simulations, except when demonstrating consequences of the absence of autocorrelated noise. *σ* controlled the degree of scan noise in the simulation and was varied across three levels of 0.8, 1.6, and 3. Before estimating activation levels, both the regressors and the simulated signal were high-pass filtered with a Gaussian weighted running line that matched the FSL method (filtering threshold 64s).

Estimation of the single trial activities was performed with both LS-A and LS-S. The LS-A and LS-S estimates were obtained by running the following GLM models for each scanner run separately:

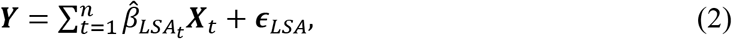

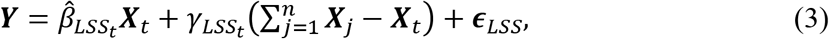

where 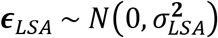 and 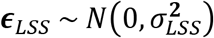. 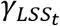 in equation 3 represents the coefficient for the nuisance regressor in LS-S method. The nuisance regressor is created by summing all the trials’ estimated BOLD activity except for that of the trial of interest. Though not reported in detail here, we also performed simulations using separate nuisance regressors for each of the trial categories as proposed by Turner et al. (2012). We found that mean centering affected both LS-S methods in a similar manner, though overall classification accuracies were higher with the separate nuisance regressors method proposed by Turner et al. (2012). To show the effect of run-wise mean centering on LS-A and LS-S estimates, we obtained run-wise mean-centered estimates by subtracting from the original estimates the mean of all the estimates within that run:

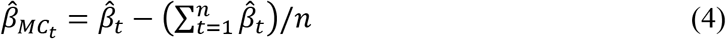

To assess the correlation between the mean estimated activation levels for trial A and trial B across runs, we simulated 2,000 runs of data (10 trials per run for each of two trial types) and estimated the single trial activity levels using LS-A or LS-S. Then the average activity levels for type A and type B were obtained by calculating the arithmetic mean of each of their respective trials’ estimates (i.e., mean of all type A’s estimates and mean of all type B’s estimates). The correlation between these 2,000 pairs of average activation levels was examined while varying several factors including ISI, presence of different types of noise (no noise, I.I.D. noise, AR noise), and presence of trial-to-trial variability in signal (e.g., *β*_*t*_ ~ *N*(3,0.5^2^); vs. *β*_*t*_ = 3).

To examine the effect of noise and ISI on cross-validated (CV) classification accuracy, we simulated 5,000 sets of data. Each set of simulated data involved three runs, each of which had 20 trials (10 trials for each of two trial types). After activation levels were estimated with each method, a leave-one-out 3-fold CV accuracy was measured by training a logistic classifier on two training runs and testing on the remaining held-out run. We repeated this procedure for each combination of ISI and scan noise.

In order to assess the effect of mean centering when there are true shifts in mean across runs, we simulated data for two different scenarios. In the first scenario, we simulated changes in activation level across runs due to attentional drifts and physiological states. We simulated 5,000 sets of data for each combination of ISI (0-4, 5-9, 10-14, 15-19, 20-24) and activity multipliers (from 0.5 to 2 in 0.1 increments), which amplified or reduced the *β*_*t*_;s. Each set consisted of 2 runs with 30 trials each (15 trials of each trial type). One run in each set was generated via normal *β*_*t*_ (i.e., formula 1), while the other run was generated with *β*_*t*_s that were multiplied by the activity multipliers. This effectively mimics a dataset where one run has elevated or depressed activity levels for both trial types due to physiological arousal, amount of attention, etc.

In the second scenario, we examined the effect of mean centering when the training and testing datasets have different trial type proportions. We simulated 5,000 sets of data for each combination of ISI (0-4, 5-9, 10-14, 15-19, 20-24) and trial type proportions (5:5, 6:4, 7:3, 8:2). For this analysis, 2 runs of data were generated in each set, with each run containing 30 trials. The two runs’ ratio of the two trial types (A and B) were complements of each other. For example, if one run contained 18 type A trials and 12 type B trials (ratio of 6:4), the other run in the iteration would contain 12 type A trials and 18 type B trials. For both scenarios, balanced CV accuracies were obtained via training a 2-fold CV design with a logistic classifier.

#### Real Data

We used the same dataset as in Mumford et al. (2012). The dataset comes from a study that examined task switching (Jimura et al., 2014) and is available online at https://openfmri.org/dataset/ds000006a. It consists of data from 14 subjects, 10 of whom had six runs and 4 of whom had five runs of data. Each run contained visual presentations of 32 plain words and 32 mirror-reversed words. The subjects’ task was to decide whether each word signified a living entity or not. Only the trials in which the subject gave an accurate response were included in the analysis, resulting in an average of 25 mirror-reversed words and 29 plain words per run. Each trial lasted 3.25 seconds, with the interquartile range of ISIs being 3.8s-7.8s. Further detail can be found in Jimura et al. (2014).

Pre-processing and activity estimations were carried out with the FMRIB Software Library (FSL, www.fmrib.ox.ac.uk/fsl). Images were motion-corrected, skull-stripped, spatially smoothed with a FWHM 5mm Gaussian Kernel, and high-pass filtered (filtering threshold of 64 sec). The design matrices were also temporally filtered. Six motion parameters were included in the GLM as nuisance variables. No temporal derivatives were included. For LS-A estimation, a separate regressor was included for each trial, whereas for LS-S estimation, a separate GLM was run for each trial with one regressor modeling the trial of interest and one nuisance regressor modeling the remaining trials. After all the single trial coefficients had been calculated, they were concatenated and registered to a standard 3mm MNI template. For classifier training, only the voxels that were inside a standard 3mm MNI brain mask were used. No feature selection or data reduction method was used.

To assess the correlation between estimated activation levels for plain and mirror-reversed words across runs, we took the median estimated activation levels for each of the two trial types in each scanner run. Because there was a total of 80 runs (across 14 subjects), each voxel had 80 median estimated activations for plain word trials and 80 median estimated activations for mirror-reversed word trials. A correlation coefficient was calculated using these 80 pairs of median activations. To show that this correlation is not entirely driven by cross-subject variance, we mean-centered the estimated activations at the subject level to remove mean differences across subjects but maintain mean differences across runs, and then re-calculated the correlation between estimated activity for plain and mirror-reversed words across the 80 runs.

Finally, to show the benefit of mean centering in out-of-sample prediction, we employed a 6-fold (5-fold for 4 subjects) leave-one-out cross-validation design to examine the out-of-sample accuracies of classifiers trained with coefficients estimated from each method. Hence a classifier was trained on 5 runs of data (4 runs for 4 subjects) and was tested on the left out run, and this process was repeated 6 times (or 5 times for 4 subjects). A linear support vector machine (SVM) was executed via MATLAB’s (www.mathworks.com) statistics and machine learning toolbox. To choose the appropriate regularization parameter, we employed a secondary leave-one-out CV with 5-fold (4-fold for 4 subjects) design. In other words, after leaving out one testing run, we employed another leave-one-out 5-fold CV only in the training set to choose the parameter combination that had the highest balanced CV prediction accuracy. Our measure of balanced accuracy was obtained by averaging the prediction accuracy of the plain word trials and that of the mirror-reversed word trials. We performed a grid search for the regularization parameter and chose the one that yielded the highest balanced CV accuracy in the training set. Then the entire training set was used to train a classifier with the selected parameter.

## Results

### Overview of the problem and the solution

Figure 1 shows six runs of simulated data and estimated activity levels that illustrate the main feature of LS-A and LS-S estimates that we are concerned with, spurious shifts in the mean estimates across scanner runs, and our proposed solution, run-wise mean centering. In these simulations, the true underlying activities for trial types A and B are independent from one another and stable across scanner runs. However, the estimated activities for trial types A and B are correlated and vary in unison across scanner runs. That is, if the estimated mean activity for trial type A is high in a run, the estimated mean activity for trial type B will also be high, and vice versa. We more fully explore the cause of this correlation below, but it is clear that there must be a source of noise in the estimates that is shared between trial types within a given scanner run. This feature of LS-A and LS-S estimates has not been previously described. Mumford et al., 2014, showed how estimates for neighboring trials within a run were correlated (positively for LS-S, negatively for LS-A), but no one has explored the potential impact of correlations across scanner runs between the estimated activity of different trial types.

As we more fully demonstrate below, these shifts in the mean activity estimates across runs hinder accurate classification, as they add systematic variability to the single trial estimates and as classifiers are trying to find a boundary between trial types that is stable across scanner runs (indeed, most cross-validation schemes train on some subset of runs and test on the left out subset). We propose a simple solution to this problem, which is to mean-center estimated trial activities within a scanner run (Fig. 1). Run-wise mean centering should better align the activity estimates across runs, thereby reducing the variance of the trial estimates when concatenating data across runs. Note that mean centering experiment-wise across all runs (i.e., subtracting out the dotted line in Fig. 1), which is a standard feature of many classification approaches, does not accomplish the same goal. This is also different from subtracting out the global univariate average activity within an ROI in order to demonstrate the existence of multivoxel pattern that is not dependent on overall activity (Esterman et al., 2009; Kamitani & Tong, 2005, Polyn et al., 2005).

**Figure 1.**
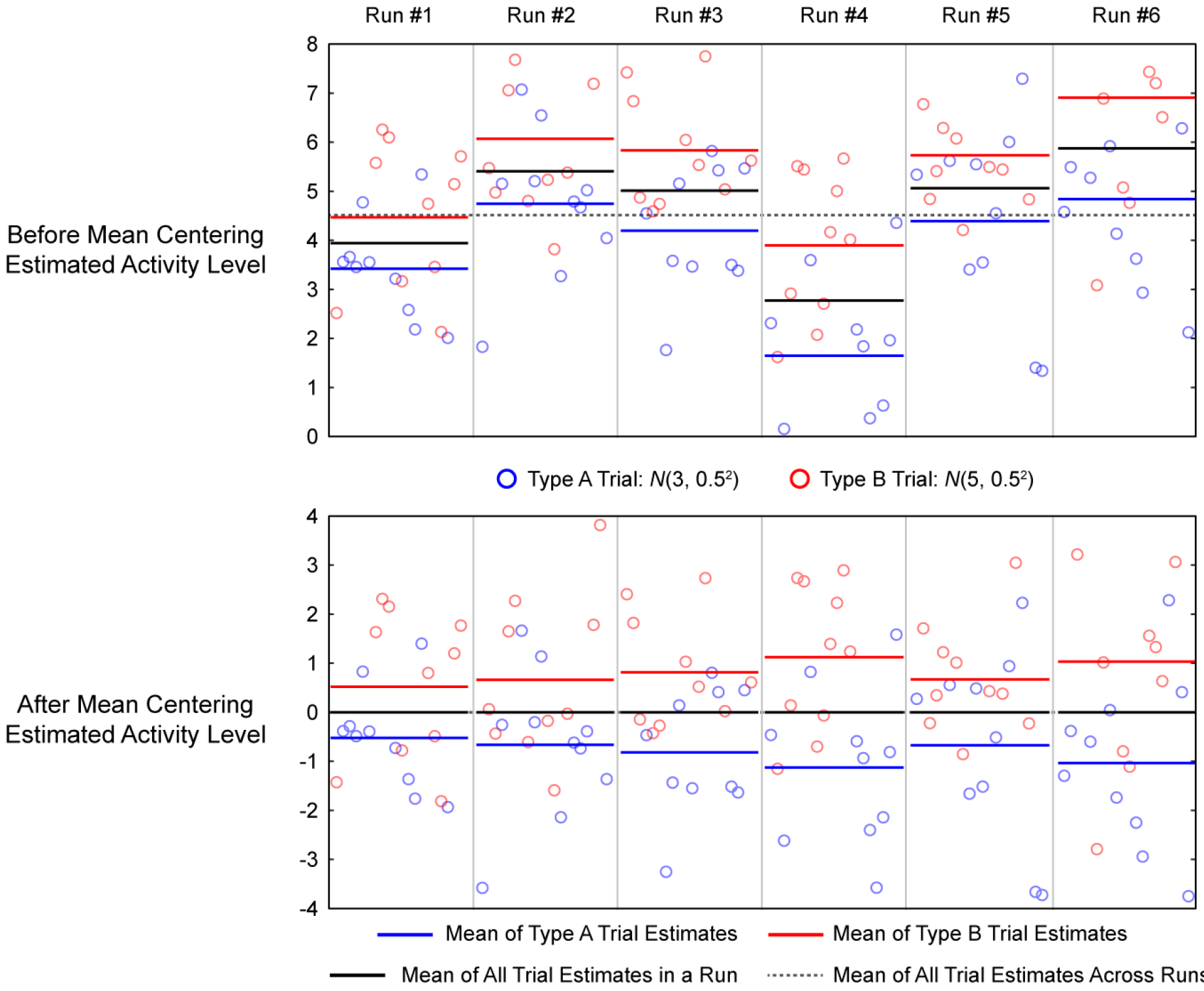
Effect of mean centering on simulated data. 6 runs of data was simulated with ISI drawn from U(0, 4), and noise std. = 0.8. Single trial estimates were performed with LS-S procedure. Top panels plot the estimated activity levels of each trial for each run. Two trial types are color-coded and their respective run-level means are also shown in color-coded bars to show spurious mean shifts. The horizontal dotted line marks the experiment-wise mean across runs. The bottom panels plot the same data but with run-level mean centering, which essentially subtracts the mean of all trial estimates in a run (shown in black bar) from all the trial estimates within the said run.

It is important to note that subtracting the mean of all trial estimates within a run will not decorrelate the means of the estimated activities of the two trial types. Rather, the means of the two trial types will be negatively correlated across runs after run-wise mean centering (bottom panel of figure 1); that is, when the mean of type A trial estimates is low, the mean of type B trial estimates is high. Hence, run-wise mean centering helps improve classification not by de-correlating the mean estimates of the two trial types, but rather by better aligning the estimates across runs. Figure 2 illustrates this and shows the histogram of the estimated single trial activity levels concatenated across 20,000 runs with and without run-wise mean centering. As expected, the distribution of single trial activity estimates for each trial type is narrower and has smaller variance after run-wise mean centering. As the difference in the means of the two distributions remains the same, this decrease in variance leads to higher separability.

**Figure 2.**
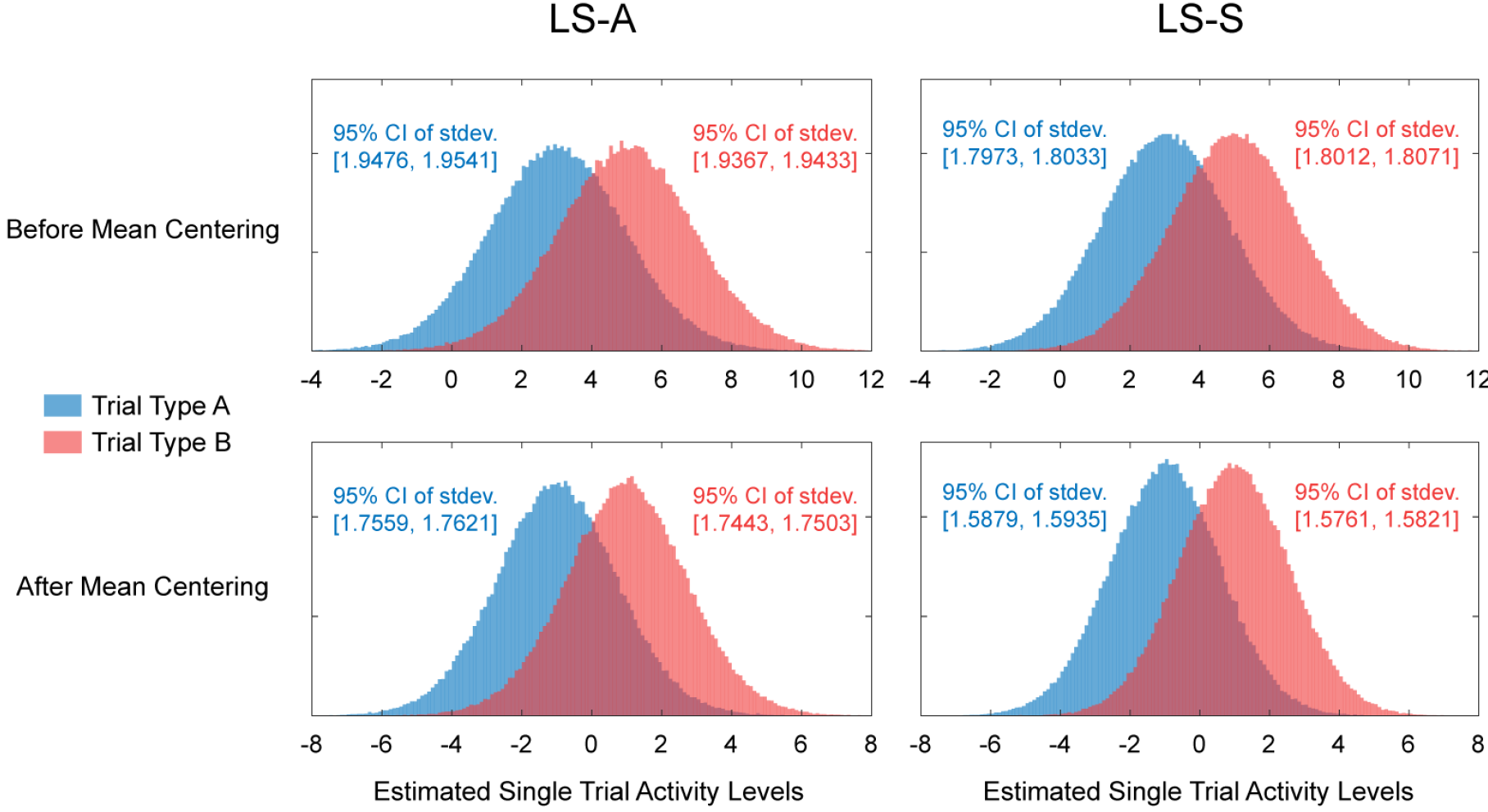
Variance of single trial estimates. Above shows histograms of estimated activity levels collapsed across 20,000 runs of simulated data using ISI ~ U(0, 4) and noise std. = 0.8. LS-A estimates are shown in the left two panels and LS-S estimates are shown in the right two panels. Top two panels show distribution of estimates without mean centering and bottom two panels show the distribution of estimates with mean centering. The bootstrap confidence intervals for the standard deviations of each of the distributions are shown in text.

### Simulation Results

To understand why the estimated activity levels of different trial types is spuriously correlated across scanner runs, we conducted a series of simulations that systematically varied the presence/absence of different potential sources of noise, both trial-to-trial variability in activity and timepoint-to-timepoint variability in scanner or measurement noise (Fig. 3). These simulations demonstrate that the estimated activations for the two trial types are correlated across runs whenever there is scanner or measurement noise. When there is no trial-to-trial variability and no noise (top two panels), LS-A perfectly returns the underlying mean activity for the two trial types, while the regularization features of LS-S, which are beneficial under realistic conditions (Mumford et al., 2012), lead to a mild level of correlation between the activity of the two trial types across runs. When trial-variability is added, this increases the variance of the activity estimates but does not systematically increase the correlation between the two trial types. Hence it seems that trial-variability in signal is not responsible for the correlation. In contrast, when noise is added to the signal, even in the absence of trial-variability, there are now strong positive correlations between the activity estimates of the two trial types across scanner runs. Similar levels of correlation are present for both LS-A and LS-S, regardless of whether the noise is (realistically) auto-correlated or (unrealistically) independent (third and fourth row panels), and regardless of whether trial-variability is present or not (fourth and fifth row panels). These simulation results show the spurious correlations occur whenever trial activity levels are estimated in the face of scanner or measurement noise.

**Figure 3.**
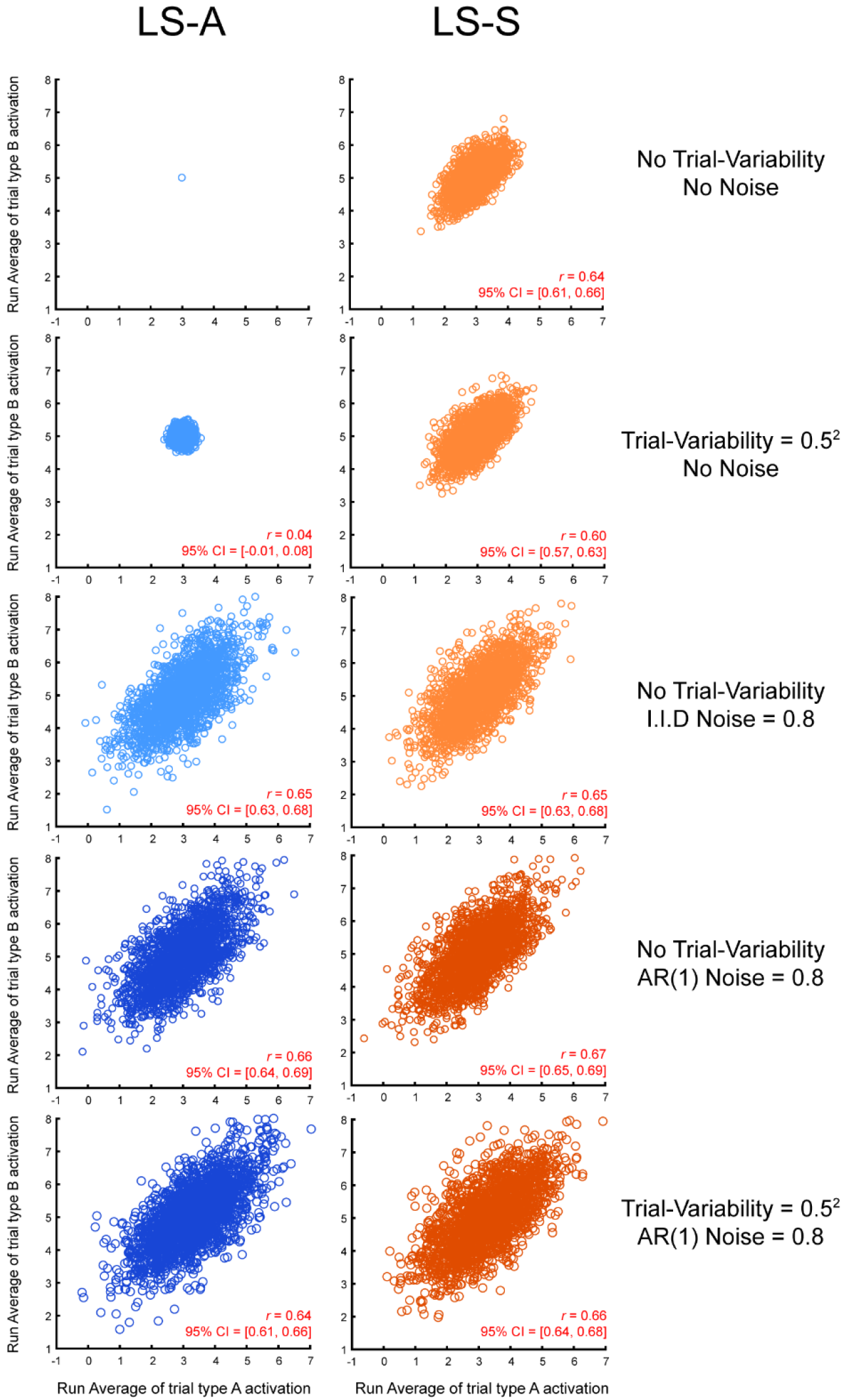
Correlated activity estimates in simulation. Correlation between the estimated activation levels of the two trial types across 2,000 simulated runs. Left panels show activation levels estimated with LS-A while the right panels show activation levels estimated with LS-S. All simulation’s ISI was uniformly drawn from U (0,4).

We also examined how this correlation varied across different ISIs (Fig. 4). The spurious correlations are strongest for short ISIs, when there is a need to deconvolve overlapping trial activities and are reduced as trials are spaced farther apart and their activity overlaps less. However, the correlation is still present even when the average ISI is 22 seconds, which is longer than most event-related fMRI designs.

**Figure 4.**
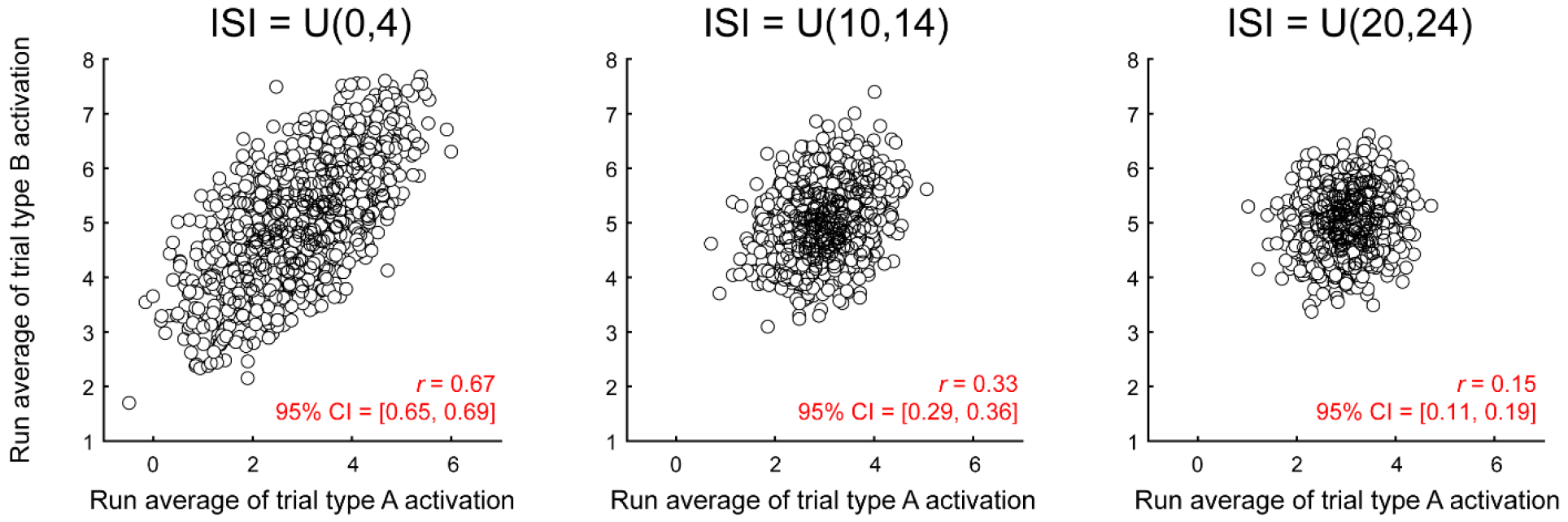
Activity estimate correlation across ISI. Correlation between the estimated activation levels of two trial types in 2,000 simulated runs, as a function of ISI, estimated with LS-S, noise std. = 0.8. The results from LS-A estimation were not much different (ISI = U(0,4): r = 0.64, 95% CI = [0.61, 0.67]; ISI = U(10,14): r = 0.36, 95% CI = [0.32, 0.40]; ISI = U(20,24): r = 0.13, 95% CI = [0.08, 0.17]).

We next turned to quantifying how much run-wise mean-centering can help out-of-sample prediction, by examining cross-validation accuracies across different inter-stimulus interval lengths and scanner noise levels. Across all tested conditions, run-wise mean-centering boosted the balanced average out-of-sample prediction accuracies (Fig. 5). Figure 5 shows boxplots of the classification accuracies for different ISI and scanner noise levels. Classification accuracies are higher with longer ISIs and with smaller scanner noise, but run-wise mean centering is helpful under all conditions. We also counted the number of simulation sets (out of 5,000) in which mean centering provided equal or better classification accuracies. When noise = 0.8, roughly 80% of all simulation sets saw equal or increased CV accuracy with mean centering, 71% when noise = 1.6, and 63% when noise = 3.

**Figure 5.**
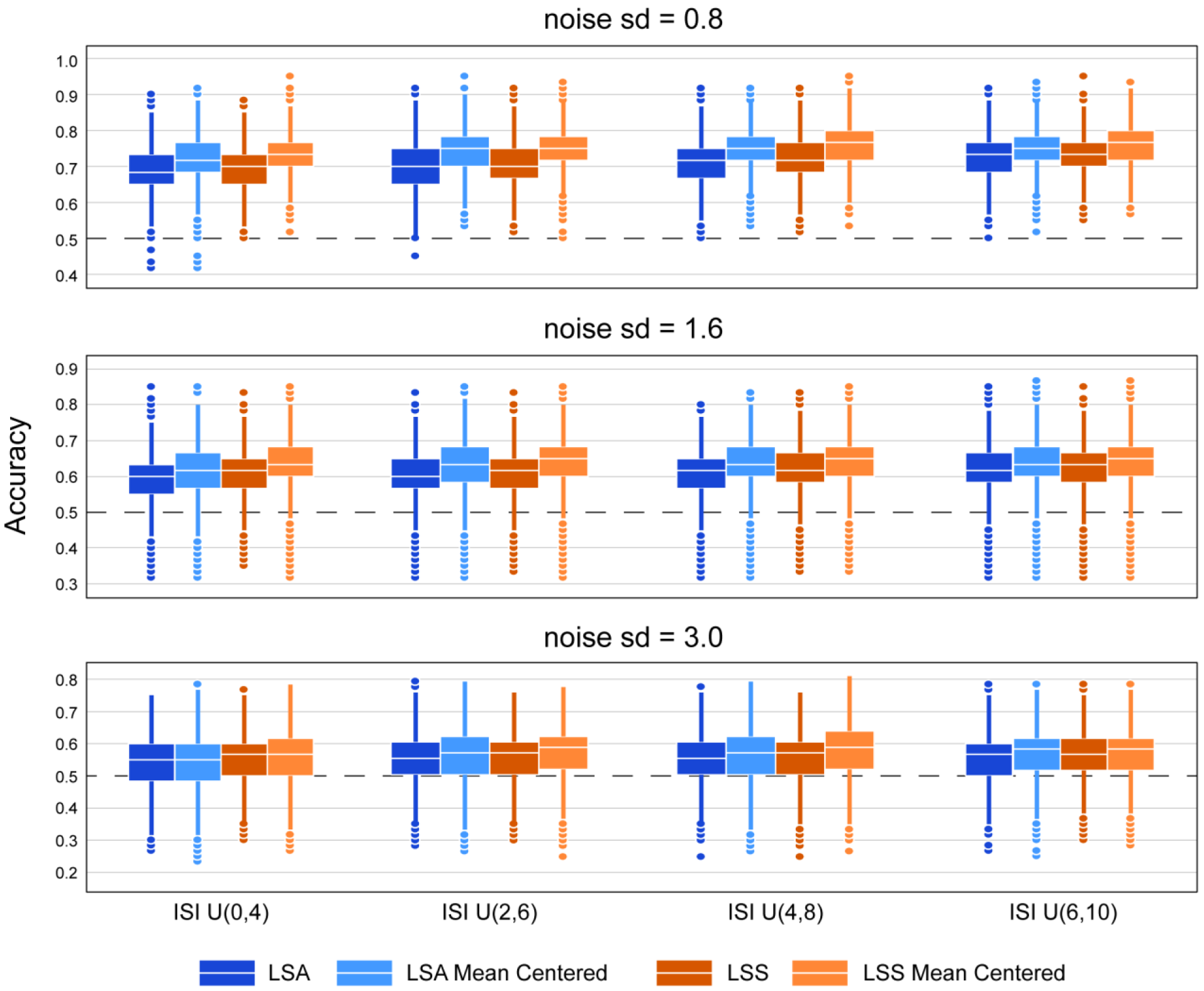
Improvement of classification accuracy in simulation. Boxplot of out-of-sample cross validation accuracies from simulated data, according to simulated ISI and noise levels and estimation method. Each boxplot shows 5,000 average CV accuracies.

These results suggest that mean centering activity estimates within a scanner run can improve multivoxel classification accuracy by correcting for spurious mean shifts imparted by deconvolution in the presence of noise. However, there are cases where we would expect there to be a *true* shift in the mean activation level between runs. We examined the effects of mean centering in two cases where true shifts in mean activation are expected, but for different reasons.

One possible reason for there to a true shift in the mean activation level between runs is fluctuation in the subject’s attentional or physiological state. Such fluctuations might act as an activation multiplier, increasing or decreasing the activity for all trial types together. However, in this case, run-wise mean centering should still prove advantageous, as the coordinated increases or decreases due to state changes across scans would add systematic noise and obscure the boundary between trial types. To illustrate this effect, we simulated training and testing datasets with different activation multipliers (Fig. 6). Regardless of mean centering, we see that longer ISIs tend to result in higher classification accuracies. More importantly however, we see that without mean centering, CV accuracy is very sensitive to the activity multiplier, decreasing in all cases when the multiplier is not exactly 1. Mean centering shows robust improvement across all combinations of ISI and activity multipliers. Interestingly, CV accuracy increases with the activity multiplier with mean centering, as mean centering allows the classifier to take advantage of the greater separation between trial types with increasing activity multipliers.

**Figure 6.**
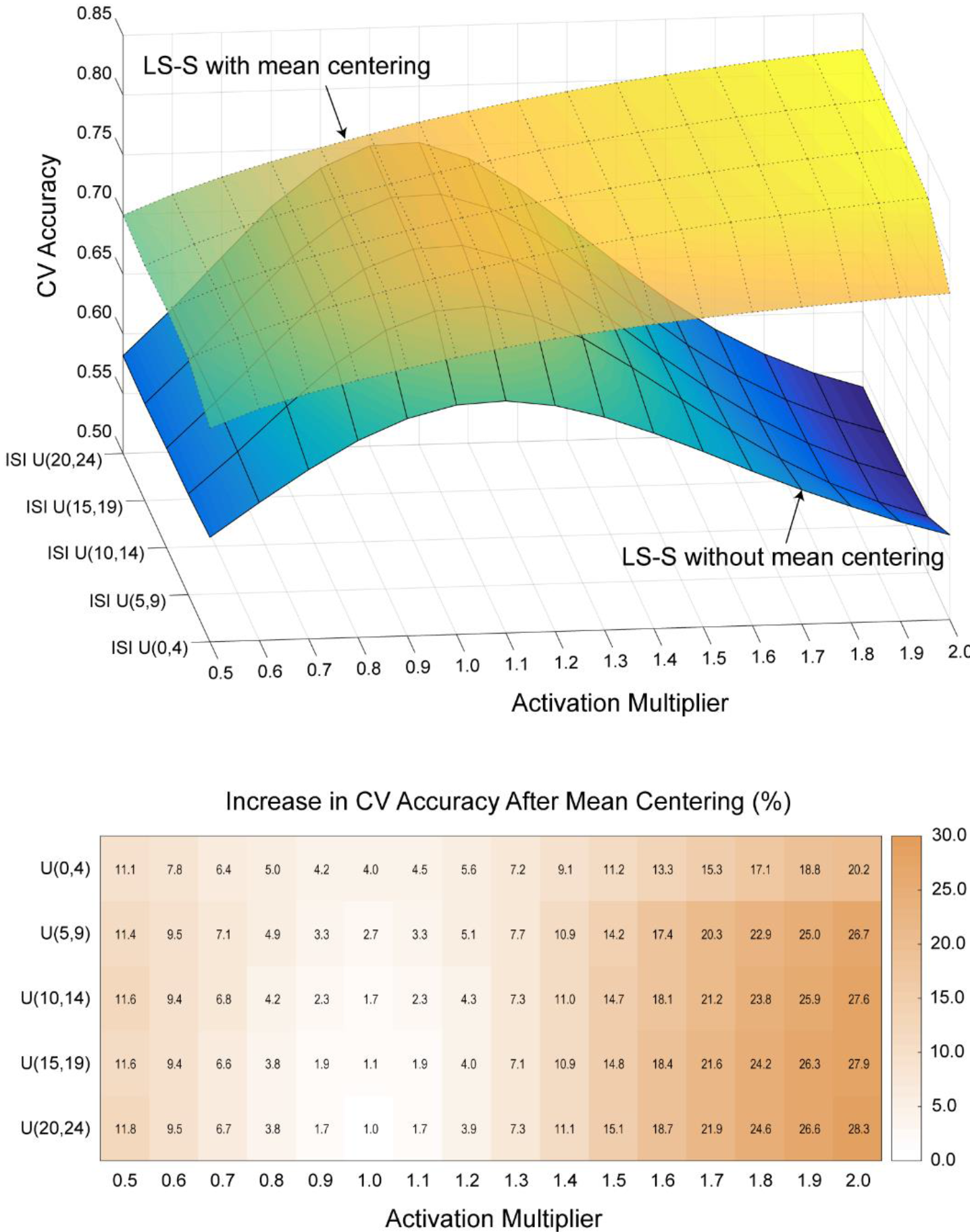
Effect of mean centering under variable global activity levels. Top panel shows the CV accuracy between two runs that have different global activity levels with and without mean centering. Bottom panel shows the difference between them. Standard errors were below 0.1% for all cells.

Another possible reason for there to be a true shift in the mean activation across runs is when runs are composed of different proportion of trials. A run with more type A trials will have an overall mean that is closer to the mean of type A trials, whereas a run with more type B trials will have an overall mean that is closer to the mean of type B trials. In this case, run-wise mean-centering should be detrimental, as subtracting out the overall mean from each run will effectively bring the activities of different trial types closer to each other and make it harder to classify. To illustrate this effect, we simulated training and testing datasets with different trial type proportions (Fig. 7). As before, we see that the benefit of run-wise mean centering is higher when the ISI is shorter. More importantly, though, we see that run-wise mean-centering only improves prediction accuracies when the proportion of trial types in the training and testing set are similar; when the proportion of trial types in the training and testing set are markedly different, imparting a true difference in the mean activation levels across the two sets, run-wise mean-centering decreases prediction accuracies.

**Figure 7.**
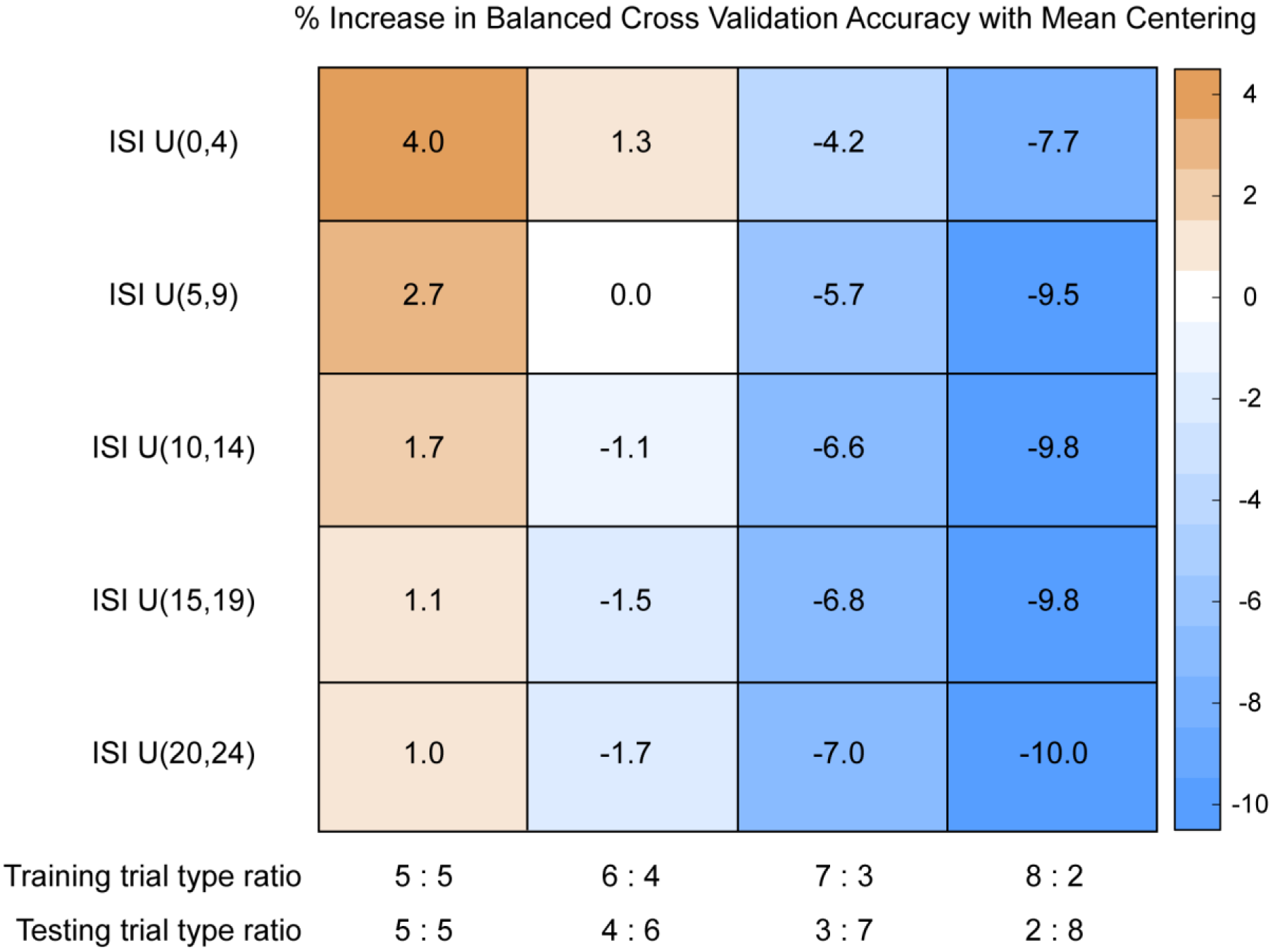
Mean-centering in unbalanced datasets. Percent increase in balanced CV prediction accuracies with mean centering. Each cell shows the CV accuracy with mean-centered LS-S coefficients minus the CV accuracy with non-mean-centered LS-S coefficients. Standard errors were below 0.1% for all cells.

### Real Data Results

To show the applicability of these concerns to real data, we used the same dataset from Jimura et al. (2014) that was used by Mumford et al. (2012). In this dataset, as in our simulations above, we observe a correlation between the estimated activation levels for plain word and mirror-reversed word trials across runs (Fig. 8; LS-A: median *r* = 0.68, LS-S: median *r* = 0.55). While our simulations considered a single hypothetical voxel, real datasets contain thousands of voxels, and so we plot the distribution of this correlation across voxels in Figure 7.

**Figure 8.**
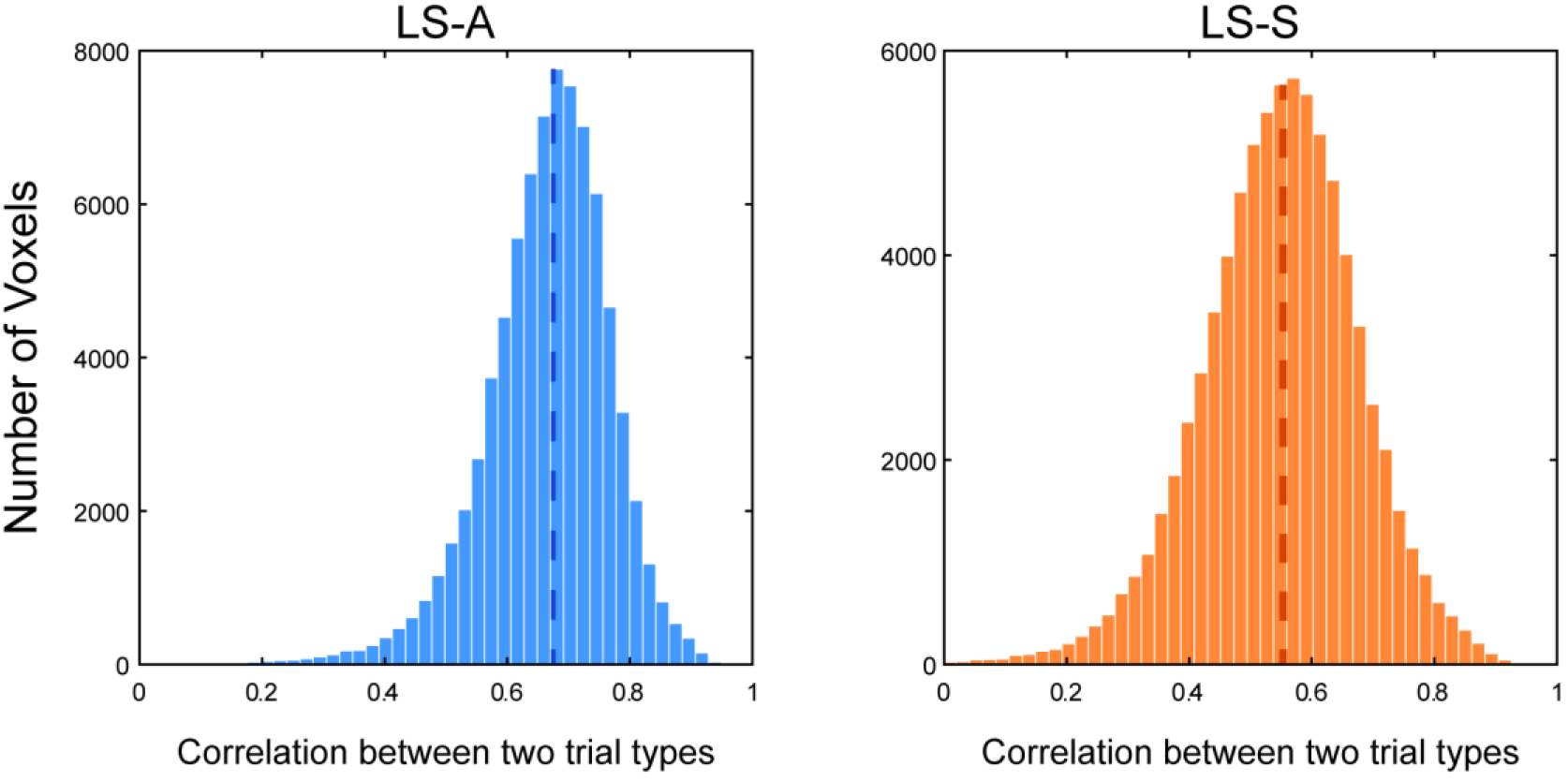
Correlated activity estimates in real data. Histogram of scanner run-level correlation between the estimated activation levels of two trial types. The left panel shows the distribution of correlation coefficients obtained via LS-A estimation, while the right panel shows that obtained via LS-S estimation. The dashed line shows the median. Both histograms are truncated at zero.

For each voxel, we calculated the correlation between 80 pairs of run-level averaged estimated activity for plain word and mirror-reversed word trial types. This combines data across all subjects, but these correlations are still robust even after removing subject-level variance, by mean centering the estimates at the subject level (LS-A: median *r* = 0.54, LS-S: median *r* = 0.39). This demonstrates that there are mean shifts in the activity estimates across scanner runs. Of course, unlike our simulations, we cannot know that these shifts are entirely spurious and imparted solely by the estimation procedure; though the proportion of trial types is the same in each run, we cannot rule out run-to-run variability in subject attentional or physiological state.

As expected given our simulation results above, though, we see a boost in out-of-sample prediction accuracies when we apply run-wise mean-centering in this dataset. For LS-A estimates, average classification accuracies increased with run-wise mean centering, though this increase was not statistically significant (Fig. 9, median difference = 2.74%, range = [−6.37% −16.41%], sign rank *Z* = 1.79, *p* = .079). For LS-S estimates, average classification accuracies significantly increased with run-wise mean centering (Fig. 9, median difference = 3.59%, range = [−1.37% −14.91%], sign rank *Z* = 2.54, *p* = .009; note that these are paired sign rank tests and that the median difference reported here does not match the difference in medians shown via boxplot).

**Figure 9.**
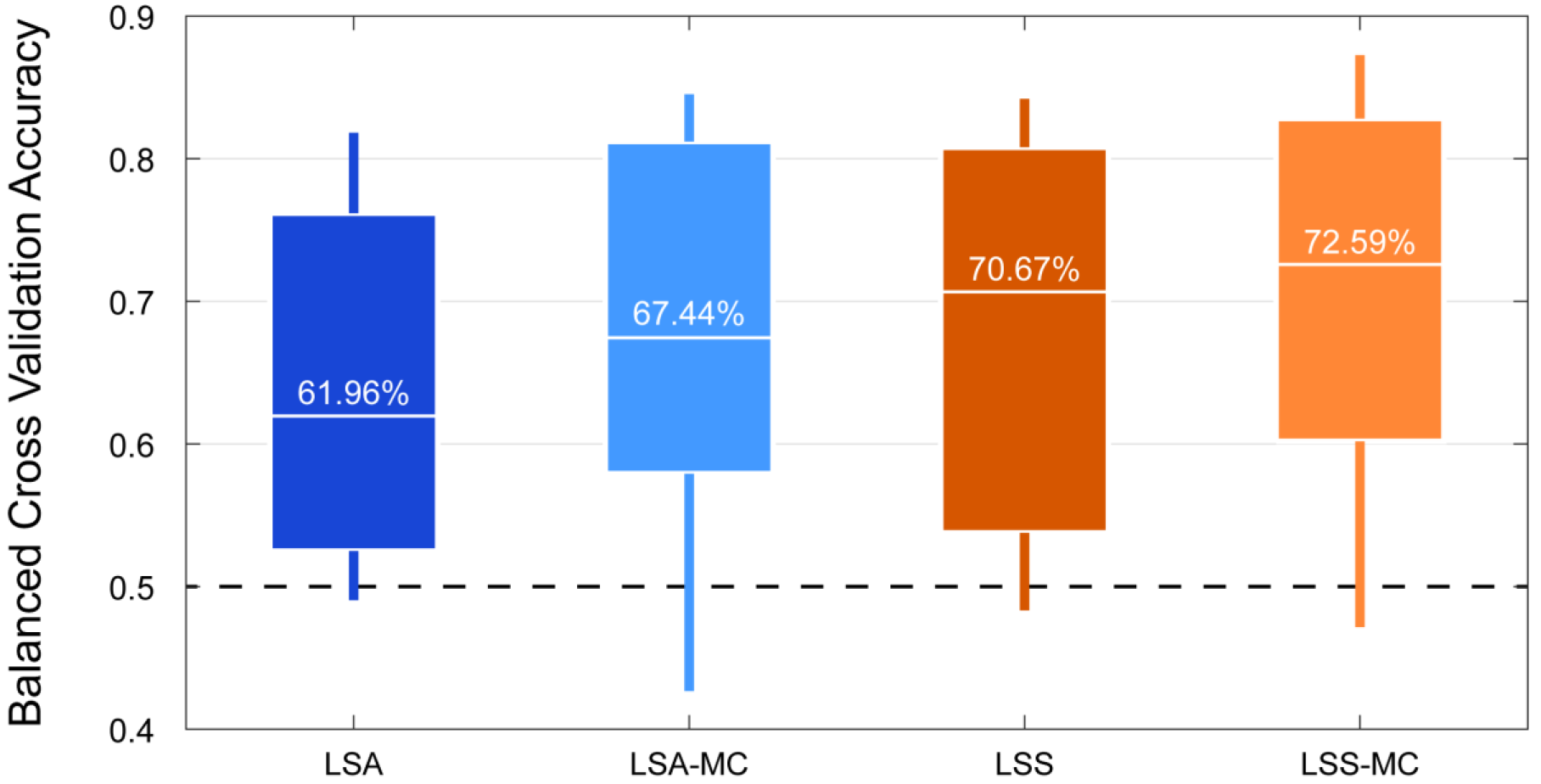
Improvement of classification accuracy in real data. Boxplot of 6 fold (5 fold for 4 subjects) out-of-sample classification accuracies from real data of 14 subjects, according to the method of estimating single trial activation levels. The box marks the interquartile range from 25% −75% while the vertical line marks the total range. The white horizontal bar and the text above it represents the median classification accuracy. MC = mean centered single trial estimates.

## Discussion

MVPA is now commonly used in fMRI research to classify mental states and/or stimuli based on neural data (Haynes & Rees, 2006; Norman et al., 2006; Pereira, Mitchell, & Botvinick, 2009). Here we show how a simple change can improve MVPA prediction accuracies across a wide range of conditions. Current techniques for estimating single-trial activations suffer from spurious correlations between the estimated activity levels of different trial types across runs, reducing classification accuracy. Mean-centering trial activation estimates within a scanner run improved MVPA classification accuracies, in both real and simulated data, regardless of estimation method (i.e., LS-A versus LS-S), and across a wide range of ISIs and signal-to-noise ratios. Furthermore, run-wise mean centering improved classification even in the presence of the kind of true shifts in mean activity across runs that might result from fluctuations in attention, fatigue, or physiological arousal. The improvements we observed in multivoxel classification accuracy (between 2% and 5% in real data) are likely to prove important in analyses that require greater statistical power, especially given that they come with practically no computational cost.

The *run-wise* mean-centering of *activity estimates* that we propose here differs from two other forms of mean-centering that are already common practice in the literature. The proposed method is to calculate the mean of all the single trial estimates within a run, and then to subtract the said mean from each of the estimates. First, it is common practice to mean-center or z-score the *raw time course* of BOLD activity before performing a GLM, and/or to include an intercept in the GLM model. Mean-centering the raw signal from a run, however, does not cause the single trial activity estimates from that run to be mean-centered. All the results in this paper are produced after mean-centering the raw BOLD activity before running a GLM. Second, many machine learning algorithms incorporate automatic scaling/standardization by default. Many machine learning algorithms and some linear models such as LASSO and ridge regression internally employ automatic z-scoring (or min-max scaling) of the predictor variables in order to aid the search for the regularization parameters and to enhance performance (e.g., Hsu, Chang, & Lin, 2003; Sarle, 1997). However, these normalizations are performed across the entire training dataset - that is, all of the scanner runs together - and the same normalization is applied to both the training and testing datasets. Such procedures remove the overall mean across all scans and all trials (the dotted line in Fig. 1), but do not correct the spurious mean shifts across runs, including between runs in the training and testing dataset. All of the MVPA analyses in Figure 9 employ this standard automatic scaling procedure, and here we see that run-wise mean-centering improves prediction accuracy over and above this default.

It is important to note some caveats to our findings. Because the benefit of run-wise mean centering comes from correcting the shifts in run-level means, it will provide no benefit if the training and testing dataset is from the same run. However, performing cross validation within the same run is generally not discouraged because it increases false positive rates (Mumford, Davis, & Poldrack, 2014). We also show that run-wise mean centering can be harmful when the ratio of trial types is markedly different between the training dataset and testing dataset. This is rarely the case in practice, as most datasets collected for MVPA typically have equal (or similar) numbers of trial types across runs by design. Nevertheless, researchers should be cautious when their research design involves discrepancies in the proportion of trial types across training and testing datasets. Future research may consider more sophisticated methods for canceling out only the spurious mean shifts that occur across runs. For example, weighted mean-centering might be considered to counteract the different proportion of trials, as long as this procedure did not require knowledge of the trial type labels in the testing dataset, as this would violate blind out-of-sample prediction. This could be workable within a Bayesian framework that uses the base rate information for different trial types.

In addition, we only considered the effect of run-wise mean-centering on MVPA classification analyses. There are other techniques, such as functional connectivity and representational similarity analyses, that also rely on single trial activity estimates. Future research should consider whether these other techniques are also affected by the noise shared across activity estimates for different trial types within the same scanner run, and whether run-wise mean-centering might also improve the sensitivity of these other techniques as well.

## Acknowledgement

This research was supported by grants from the National Institute of Mental Health [R01-MH098899] (to Kable and Gold). We thank Chris Glaze for his insights on this research.

